# Isolation and maintenance of *Batrachochytrium salamandrivorans* cultures

**DOI:** 10.1101/796920

**Authors:** Kristyn A. Robinson, Kenzie E. Pereira, Molly C. Bletz, Edward Davis Carter, Matthew J. Gray, Jonah Piovia-Scott, John M. Romansic, Douglas C. Woodhams, Lillian Fritz-Laylin

## Abstract

Discovered in 2013, *Batrachochytrium salamandrivorans* (*Bsal*) is an emerging amphibian pathogen that causes ulcerative skin lesions and multifocal erosion. A closely related pathogen, *Batrachochytrium dendrobatidis* (*Bd*), has devastated amphibian populations worldwide, suggesting that *Bsal* poses a significant threat to global salamander biodiversity. To expedite research into this emerging threat, we seek to standardize protocols across the field so that results of laboratory studies are reproducible and comparable. We have collated data and experience from multiple labs to standardize culturing practices of *Bsal*. Here we outline common culture practices including a media for optimal *Bsal* growth, standard culture protocols, and a method for isolating *Bsal* from infected tissue.

## 1. Introduction

Two species of chytrid fungus, *Batrachochytrium dendrobatidis* (*Bd*)(Longcore et al. 1999) and *Batrachochytrium salamandrivorans* (*Bsal*) (Martel et al. 2013), are the etiological agents of chytridiomycosis, a necrotic skin disease that is a major driver of the global decline in amphibian biodiversity (Scheele et al. 2019).

*Batrachochytrium dendrobatidis (Bd)* was the first pathogen known to cause chytridiomycosis. Its isolation in 1999 (Longcore et al. 1999) prompted vigorous efforts to understand *Bd*’s basic biology, geographic distribution, and the major factors promoting its spread and persistence (Berger et al. 1998, Woodhams & Alford 2005, Lips 2016). *Bd* can infect a broad range of amphibian hosts, including frogs (Scheele et al. 2019), salamanders (Chatfield et al. 2012), and caecilians (Gower et al. 2013). *Bd* has been detected on every amphibian-inhabited continent and is presumed to have spread through the globalized trade of infected amphibians originating from endemic areas (Schloegel et al. 2012). The impact of *Bd* has been particularly pronounced for frogs, and has already driven 90 species to extinction (Kilpatrick et al. 2010, Scheele et al. 2019). Although many questions remain unanswered, the ability to isolate *Bd* from amphibians and maintain *Bd* cultures in the laboratory (Garner et al. 2016, Waddle et al. 2018, Cook et al. 2018) has advanced the understanding of its epidemiology and enabled the evaluation of mitigation strategies to prevent further declines and extinctions (Waddle et al. 2018).

*Batrachochytrium salamandrivorans* (*Bsal*) was identified in 2013 following a sudden crash in Fire Salamander (*Salamandra salamandra*) populations in the Netherlands (Martel et al. 2013). While both frogs and salamanders can be infected by experimental exposure to *Bsal* (Stegen et al. 2017), only post metamorphic salamanders appear to develop ulcerative skin lesions and multifocal erosion (Van Rooij et al. 2015). Non-Asian salamanders belonging to the family Salamandridae are especially susceptible to *Bsal* and often experience high levels of disease and mortality (Martel et al. 2013).

*Bsal* is thought to have originated and naturally coexist in amphibian communities throughout Asia without causing apparent harm (Laking et al. 2017). Like *Bd, Bsal* is predicted to continue to spread by way of the pet trade of amphibians carrying subclinical *Bsal* infections (Yuan et al. 2018, Sabino-Pinto et al. 2018). Though *Bsal* chytridiomycosis outbreaks have not yet been observed outside of Europe, *Bsal* poses a significant threat to global salamander biodiversity and is predicted to soon spread to North America, if it has not already(Watts et al. 2019).

To prevent a *Bsal* pandemic that could rapidly drive salamanders to extinction, we need to develop conservation and disease management strategies that are based on controlled laboratory research. This requires established techniques for maintaining laboratory cultures of *Bsal*, and there is growing interest in a common set of protocols to standardize practice among laboratories. Here, we provide an overview of maintenance of *in-vitro Bsal* cultures and describe recommended protocols for: 1) culturing *Bsal* in liquid and solid media, 2) isolating specific *Bsal* life stages, and 3) extracting *Bsal* from infected tissue.

## 2. Basic techniques for culturing *Batrachochytrium salamandrivorans* (*Bsal*)

### 2.1 Biosafety

*Bsal* is an amphibian-specific pathogen and does not pose a significant risk to humans. To prevent the spill-over of *Bsal* outside of the laboratory, it is imperative that high levels of biosafety (at least the equivalent of United States Biosafety Level 2) are observed when working with live cultures. This includes conducting all culture work in a biosafety cabinet. The following resources may be useful for working with your institution to develop a detailed biosafety protocol: (Burnett et al. 2009) outlines general biosafety practices when working with pathogenic microorganisms, (Van Rooij et al. 2017) reviews chemical disinfectants and necessary exposure times for killing *Bsal,* and an example protocol for approval by institutional biosafety committees can be found at salamanderfungus.org. This example protocol includes recommendations for containment and disinfection approved by USDA, although *Batrachochytrium* and other wildlife pathogens are not currently under the purview of the USDA Animal and Plant Health Inspection Service (website: http://www.aphis.usda.gov/).

### 2.2 Life Cycle

Like *Bd* and other zoosporic fungi, the life cycle of *Bsal* is characterized by a free-swimming infective stage known as a zoospore and stationary reproductive stage called a sporangium (**Figure 1**). Though population genetic evidence suggests that *Bd* may be capable of both sexual and asexual reproduction (Morgan et al. 2007), the production of asexual zoospores is presumably the primary reproductive mode for both *Bd* and *Bsal. In vitro,* zoospores give rise to thalli containing one (monocentric) or multiple (colonial) sporangia. *Bsal* sporangia have been reported to produce two types of infective spores, the motile zoospore described above, and a buoyant encysted spore that, in the wild, is hypothesized to float to the surface of aquatic habitats and aid in transmission (Martel et al. 2013, Stegen et al. 2017).

**Figure 1.**
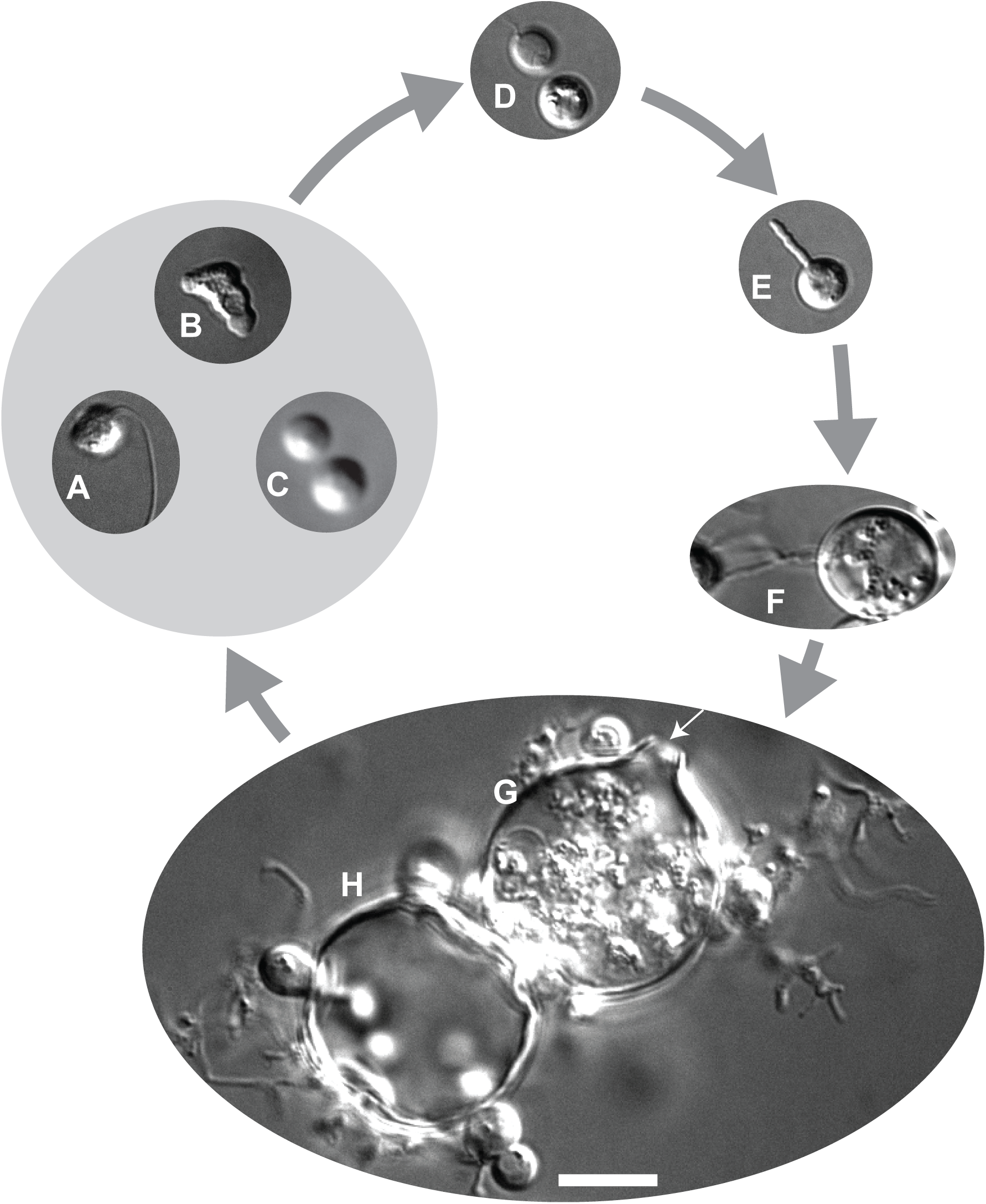
*In vitro* life cycle of *Bsal*. Two types of zoospores have been observed. **A+B.** Motile zoospores can swim through their environment using a posterior flagellum and amoeboid crawl on surfaces. **C.** Encysted zoospores have a cell wall and float in media. **D.** Once a spore is ready to colonize, it will encyst and retract its flagellum. **E.** Cysts develop one or several rhizoids. **F.** Thalli (immature sporangia) increase in volume before maturation. **G.** The mature sporangia produce zoospores internally and assemble a discharge tube (indicated by arrow). **H.** Zoospores are then released from the sporangia via dis-charge tube and begin the cycle again. Cells were imaged using a 40X air (1C) and 100X oil objective (all others). Scale bar represents 10 µm.

### 2.3 Media

*Bsal* grows well in both liquid and solid media (see ‘Recipes’). While liquid media results in more uniform growth and better sporulation, solid media allows for the formation of colonies that are not easily dislodged from the agar surface, which can be advantageous for some applications. Solid media is also best for shipping *Bsal* cultures because of the reduced potential for contamination.

It is important to carefully consider what kind of medium to use as this decision influences *Bsal* growth (see “Recipes”). To determine which is optimal, we estimated growth rates of *Bsal* zoospores grown in a variety of liquid media types by tracking the change in optical density (**Figure 2**). While all tested media facilitated *Bsal* growth, the growth rates were highest in TGhL and Tryptone media compared to potential alternatives (Tukey, p < 0.01). Moreover, the growth rates of *Bsal* grown in half-strength TGhL were significantly higher compared to cultures grown in 1% Tryptone media at either full or half-strength (**Figure 2**). Based on these data, and a desire to promote uniformity in culturing across laboratories, we recommend using half-strength TGhL media to maintain *Bsal* cultures.

**Figure 2.**
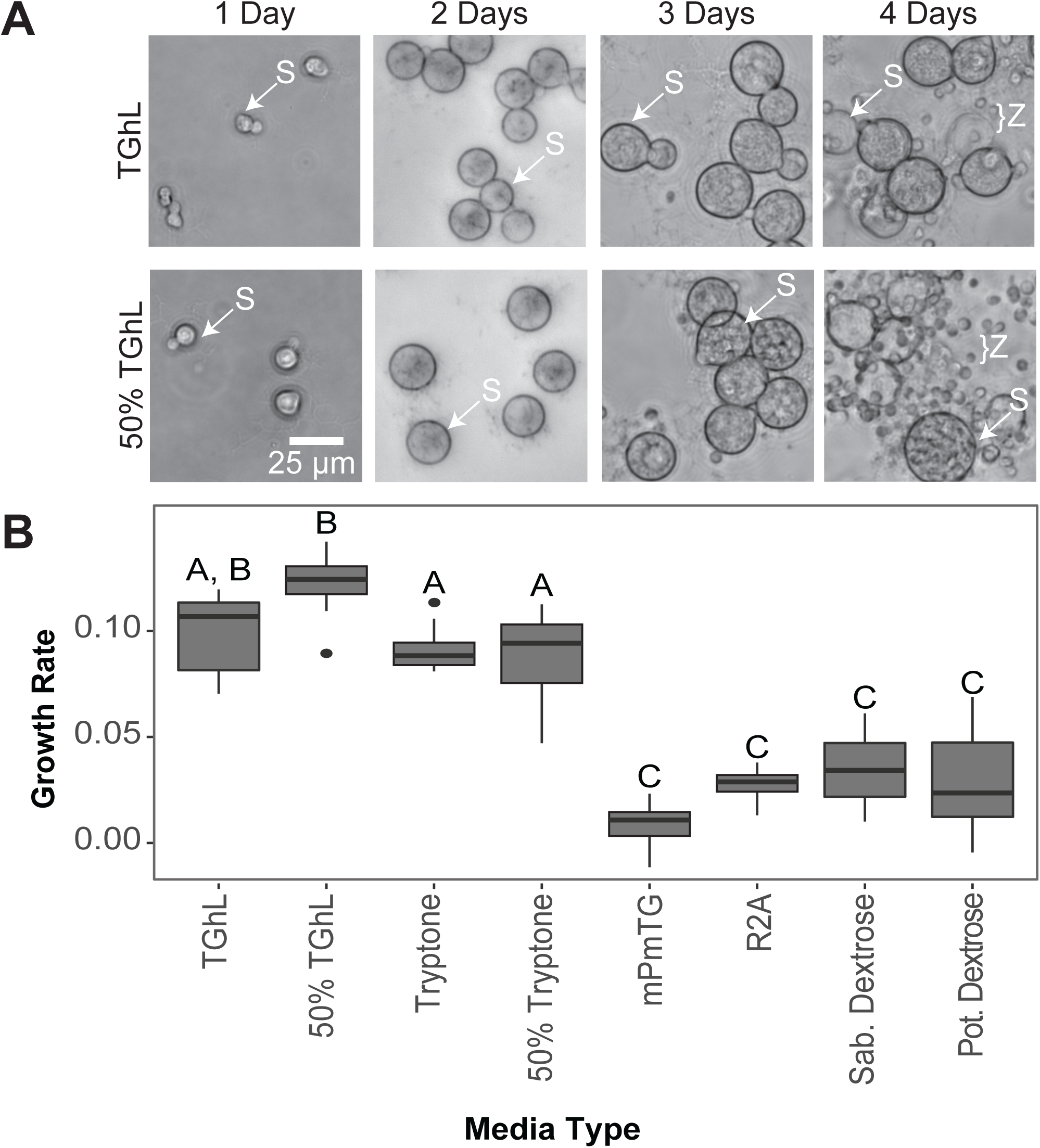
*Bsal* growth in different media types. **A.** *Bsal* was grown in half (50%) and full-strength TGhL liquid media for four days. Most sporangia (S) grown in half-strength TGhL release zoospores (Z) approximately 24 hours earlier than sporangia grown in full-strength TGhL. **B.** Growth in different media types was compared using a growth assay. *Bsal* zoospores (5e4 per well) were inoculated into the wells of a 96-well TC treated plate containing the indicated media. Growth was measured daily by optical density (OD) using an Omega FluorStar spectrophotometer at 492 nm for five days. Growth rate was calculated as the slope of OD measurements through time. Growth rate significantly differed among medias (ANOVA, F_8,62_ = 54.98, p < 0.001). Treatments with different letters are statistically significant from each other, but treatments with the same letter are not.

### 2.4 Antibiotics

While practicing good sterile technique should minimize fungal and bacterial contamination, some laboratories choose to include ampicillin and /or streptomycin in the culture media (see ‘Antibiotic Recipes’). Although the use of antibiotics may provide additional protection against microbial contamination, antibiotics must be used with caution because of their potential to alter gene expression and regulation of cultured cells (Ryu et al 2017). We recommend that the experimental needs of the lab be carefully evaluated when deciding whether to supplement media with antibiotics.

### 2.5 Growth temperature

*Bsal* grows well at 15 C and can be stored for weeks to months at 4 C. *Bsal* does not tolerate warmer temperatures (thermal maximum = 25 C) and dies rapidly if left at room temperature for an extended period of time (Martel et al. 2013). We recommend purchasing an incubator capable of holding a steady below-room temperature before beginning to culture *Bsal*. Despite the thermal limitations of *Bsal*, proper biosafety procedures must be followed prior to disposing of *Bsal* cultures and generated wastes (see ‘Biosafety’).

## 3. Subculturing

Subculturing, also known as passaging, is the addition of cells from a previous culture to fresh media (liquid or solid) to generate a new culture. We recommend regular subculturing to maintain uniform and reproducible cultures. Regular subculturing also reduces variability between experiments by minimizing differences due to aging sporangia, depletion of nutrients, and the buildup of cell waste and cell debris. We recommend recording and including passage numbers in publications as there is evidence that *Bd* can lose virulence after multiple passages in culture (Langhammer et al. 2013, Refsnider et al. 2015, Lips 2016), and we suspect that the same is true for *Bsal*.

### 3.1 Subculturing in liquid media

To subculture *Bsal,* an aliquot of mature culture is added to a clean sterile flask containing fresh media that has been pre-chilled to 15 C. The new culture flask is moved to the 15 C incubator and left to grow. Cultures inoculated with zoospores grow more reproducibly than cultures started from mixed stage cultures containing both zoospores and sporangia. Because tissue culture (TC) treated flasks make it easy to separate zoospores that are suspended in liquid media from sporangia which adhere tightly to the flask walls, we recommend growing *Bsal* in TC treated plug sealed flasks.

Growing cultures at high densities can cause an accumulation of cell waste and deplete nutrients that may limit cell growth. Growth of *Bsal* at very high densities can result in an irrecoverable state of arrested growth characterized by sporangia that fail to mature and sporulate. To maintain high, but not overcrowded, cell densities, we routinely start liquid cultures using both 1:10 (1-part zoospore culture from old flask, to 9 parts fresh media) and 1:20 ratios to ensure robust growth in at least one flask of cells. This technique results in a final concentration of around 3e4 zoospores per mL without having to count zoospores at each passage. To prevent the overgrowth of cultures, we recommend subculturing on a regular schedule. For example, growing at 15 C with half-strength TGhL, we subculture at 1:20 ratio every four days. We also recommend storing the old culture flask at 4 C for one to two weeks as a backup in case the new culture becomes contaminated.

### 3.2 Synchronizing liquid cultures

A synchronous culture contains cells in the same growth stage and can be quite useful for experiments that require large numbers of zoospores. To fully synchronize a culture, we recommend adding 6 to 7e6 age-matched zoospores (see ‘Harvesting Zoospores’ and ‘Counting zoospores’ sections below) suspended in half-strength TGhL liquid media to a 75 mm^2^ TC treated flask. Typically, a newly synchronized *Bsal* culture grown in this way will release zoospores four to six days later (see ‘Harvesting Zoospores’). In our experience, adding too many or too few zoospores results in a less synchronous culture.

### 3.3 Subculturing on solid media

*Bsal* grows well on solid media made of full or half-strength TGhL or 1% Tryptone agar with or without added antibiotics (see ‘Antibiotic Recipes’). While it is possible to subculture *Bsal* on solid media using cells grown on a plate (plate to plate subculturing), we recommend subculturing on solid media using liquid-grown cells (liquid to plate subculturing). We typically add ∼5e6 total zoospores harvested from liquid culture (see ‘Harvesting Zoospores’ and ‘Counting zoospores’) to each agar plate pre-chilled to 15 C. Sterilized glass beads or glass spreaders are then used to spread the zoospores evenly across the plate.

Mixed stage cultures can also be used to inoculate solid media and can be useful for rapidly producing large numbers of zoospores. First, use a sterile cell scraper to dislodge sporangia from the wall of the TC treated flask and then gently swirl the flask to homogenize the culture. Add 1 mL of the mixed culture to a pre-chilled agar plate and spread by tilting the plate side to side.

Once an agar plate has been inoculated with zoospores or a mixed stage culture, place plates agar side down at 15 C to allow the added liquid to completely soak into the agar (approximately 30 minutes). To determine whether the liquid has been sufficiently absorbed, agar plates are observed while being tilted side to side. If a liquid sheen is present on the surface of the plate but no moving liquid is observed then the plate is wrapped in parafilm, turned with the agar side up, and stored at 15 C for the remainder of the growth period. If liquid freely moves across the plate, the liquid is given additional time to absorb. Grow until obvious colonies develop and zoospores can be seen using a microscope (**Figure 3**). Because *Bsal* is sensitive to desiccation, agar plates should be carefully monitored to ensure they do not dry out. If plates do dry out, a humid chamber can be used to store plates in the incubator; a clean baking pan covered with aluminum foil along with a beaker of sterile water works well.

**Figure 3.**
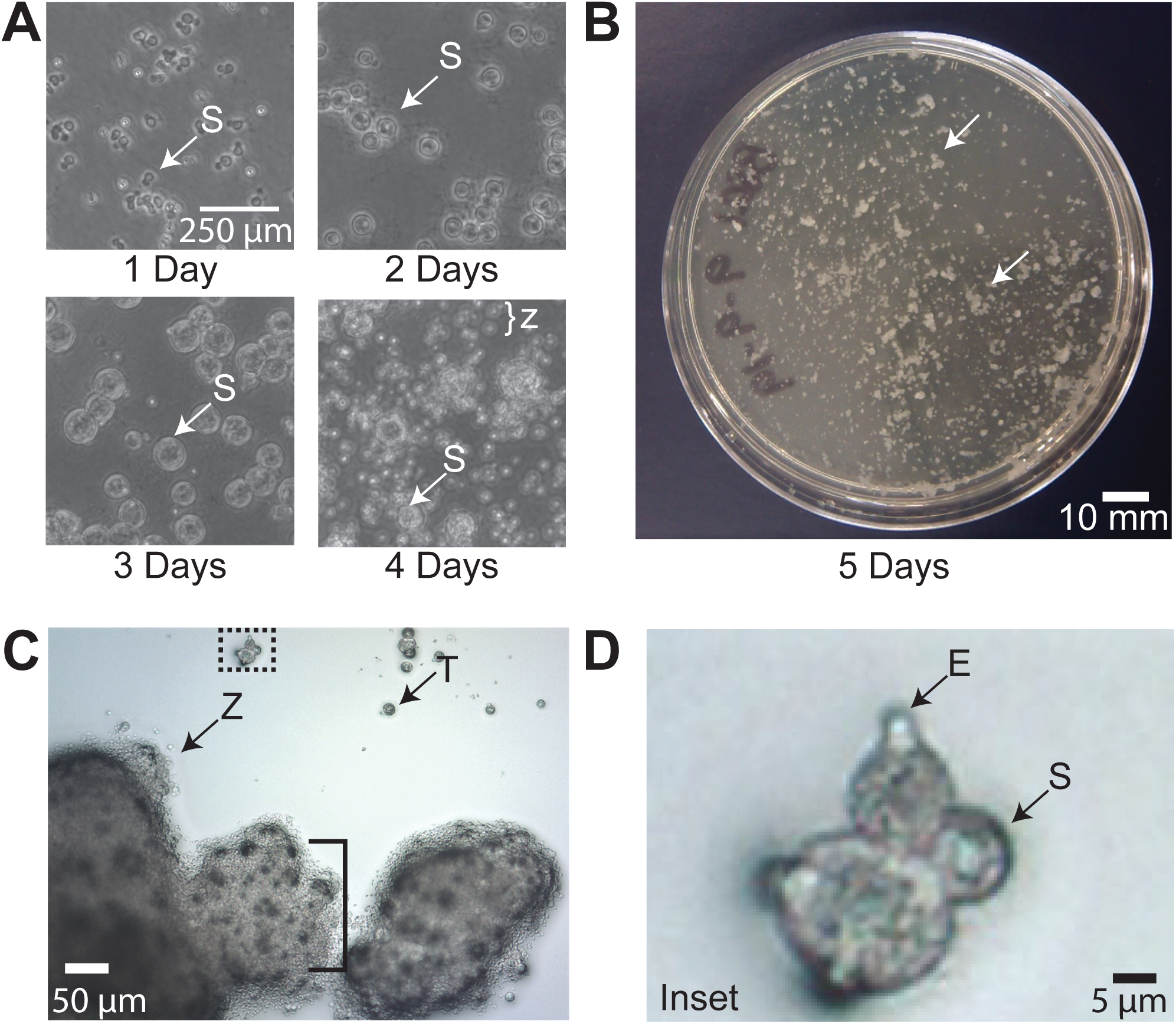
*Bsal* growth on solid media agar plates. **A.** Zoospores (Z) were inoculated on half-strength TGhL plates with 1% agar and no antibiotics. Sporulation begins after three days of growth with robust zoospore release from sporangia (S) after four days. **B.** Growth on full-strength TGhL solid media with antibiotics after four days. This plate was inoculated with a mixed stage culture grown in a TC treated flask for five days. Colonies appear as translucent to white smooth aggregations – examples are indicated by white arrows. **C.** *Bsal* colonies grown on full-strength TGhL solid media with antibiotics after a four-day incubation. Large gray structures represent aggregations (bracket) of maturing sporangia. Zoospores appear as small translucent spherical structures and are abundant at the perimeter of these aggregations, while larger spherical structures represent thalli (immature sporangia; T) and sporangia. **D.** An inset of C as indicated by the dotted box. A discharge/exit tube (E) appears as a semi-translucent growth off the side of a sporangium.

### 3.4 Handling contamination

Proper sterile technique prevents most incidences of contamination (Coté 2001). Over the course of a culture’s growth period, periodic gross and microscopic observations should be performed to verify that cultures remain healthy and uncontaminated. If available, we recommend using an inverted microscope which allows daily observation of the culture directly in the flask. It is worth noting that *Bsal* can develop biofilm-like aggregates when grown at high cell densities. Bacterial cells can be easily differentiated from *Bsal* based on differences in size (bacterial cell diameter < 2 um; zoospore diameter 4 to 5.5 um; sporangia diameter < 15.7 um), and morphology. *Bsal* zoospores often develop germ tubes (Martel et al. 2013) and can adopt an ameboid morphology similar to *Bd* (Longcore et al. 1999, Fritz-Laylin et al. 2017) that when observed under a microscope might be mistakenly identified as a biological contaminant. If contamination is suspected and cannot be ruled out via microscopic examination, subculture to solid media. Contamination of cultures grown on solid media is more easily determined because of the morphological disparities between *Bsal* colonies and those of other non-chytrid microbes (**Figure 3**).

If a culture becomes contaminated, we recommend immediately disposing of the infected culture. Clearing *Bsal* from contaminants is time consuming and often unsuccessful. New cultures can be restarted from cultures stored at 4 C that show no signs of contamination or from a frozen stock (see ‘Cryopreservation’).

## 4. Harvesting Zoospores

A synchronized liquid culture of *Bsal* can yield a vast quantity of zoospores from a single flask, with concentrations up to 2 to 8e6 zoospores per mL. Therefore, we recommend that *Bsal* cultures grown in liquid culture media are first synchronized prior to attempting to harvest zoospores (see ‘Synchronizing liquid cultures’). If a synchronized culture is not available, zoospores can be harvested from solid media (see ‘Subculturing to solid media’ and ‘Harvesting zoospores from solid media’). To determine whether a synchronized *Bsal* culture is suitable for harvesting, examine it under a microscope. If sporangia have developed discharge tube caps or papillae (**Figure 1**), and have released a few zoospores, the culture can be used for harvesting.

### 4.1 General harvesting from liquid culture

This method is used for harvesting large numbers of zoospores. To begin, remove half-strength TGhL media from the flask and replace with 10 mL of sterile Bonner’s Salts, mPmTG, or fresh half-strength media (see ‘Recipes’), depending on the subsequent use of the zoospores. The flask is then returned to 15 C for five hours to overnight to allow zoospores to collect in the chosen liquid. After the incubation period, check for zoospore release using a microscope. If zoospores are present, collect the liquid media from the flask and, if desired, filter to remove any remaining sporangia (see ‘Filtering zoospores’). Determine the concentration of zoospores using a hemocytometer (see ‘Counting zoospores’), and use for downstream applications, such as exposure experiments, growth inhibition assays, and/or subculturing.

### 4.2 Age-matched harvesting from liquid culture

For some applications, it can be helpful to start with a precisely age-matched population of zoospores. This protocol is similar to the general harvesting outlined above but uses a much shorter collection time. Start with a synchronized culture grown in a TC treated flask; when the sporangia in the culture are just beginning to release zoospores, pour out the media from the side of the flask opposite to adhered sporangia to prevent dislodging sporangia from the flask wall. Rinse the remaining zoospores from the flask by adding 10 mL of fresh media by flowing it gently across the sporangia and out of the flask. Sterile Bonner’s Salts, mPmTG, or fresh half-strength media (see ‘Recipes’) can be used for washing based on the intended use of the zoospores. Repeat this wash step a total of three times, being careful to not allow cells to dry out. After the final wash, add 10 mL of fresh liquid (the same used for the wash steps) to the flask, and check to ensure that nearly all zoospores have been removed. Only a few swimming zoospores should be visible when the flask is examined under an inverted microscope.

Return the flask to 15 C for 2 hours to allow for fresh zoospores to be released. The specific incubation time depends on how closely age-matched the population needs to be; in general, longer incubations result in a higher number of less tightly age-matched zoospores. After incubating, collect the newly released zoospores and, if necessary, centrifuge at 2500 relative centrifugal force (rcf) for 5 minutes to pellet cells and resuspend for desired zoospore concentration.

### 4.3 Harvesting zoospores from solid media

Zoospores can also be harvested from *Bsal* grown on solid media (see ‘Subculturing on solid media’). Typically, plates are ready for harvesting 5 to 6 days after inoculation. To determine whether an agar plate is ready for harvesting, examine it under a microscope (inverted or compound) turned with agar side up. If large clumps of sporangia and active swimming zoospores are visible using a 10X objective (**Figure 3**), the plate can be used for zoospore harvesting.

To stimulate zoospore release from sporangia, flood the plate with 1 to 2 mL of liquid media and incubate at 15 C for 30 minutes. TGhL media, Nanopure water, Provosoli medium and Bonner’s Salts pre-chilled to 15 C (see ‘Recipes’) are all acceptable for flooding plates, depending on the intended use of the zoospores. After 30 minutes, check the plate to see if sufficient zoospores have been released. If so, add an additional 1 mL of the chosen medium to the plate, pipetting the liquid over the plate several times before transferring into a sterile tube. Multiple rounds of plate flooding and collection can maximize zoospore collection from the same plate, although this may dilute the sample. Collected zoospores can be centrifuged at 2500 rcf for 5 minutes and resuspended in the appropriate volume of media or buffer for downstream applications. To remove remaining sporangia, the resuspended zoospores can be filtered (see ‘Filtering zoospores’).

### 4.4 Filtering zoospores

After harvesting zoospores from liquid or solid media, some laboratories also filter zoospores to further remove thalli and sporangia, particularly inoculum for animal challenge experiments (e.g. Carter et al 2019). The collected zoospores are passed through a Büchner funnel lined with Whatman filter paper or 20 μm nylon mesh (previously autoclaved). Because Whatman filter paper can absorb small volumes of liquid media along with zoospores, we recommend using nylon mesh for filtering volumes less than 2 mL.

### 4.5 Counting zoospores

For many experiments, including estimating lethal-dose 50 concentrations, it is helpful to start with a known concentration or number of cells. The most reproducible way to count cells is to count zoospores, as each zoospore represents one colony-forming unit, while sporangia can, in principle, give rise to many colonies. We recommend using a hemocytometer (for introduction to hemocytometer use, see (Absher 1973)) and a 40X objective. Zoospores swim quickly and make accurate counting difficult. Therefore, we typically fix zoospores prior to counting using Lugol’s iodine (1:100 dilution) that stains cell membranes, allowing greater contrast under the microscope. Fixation also reduces the likelihood of contaminating lab surfaces with live zoospore cultures. For studies where zoospore concentration is important, concentration can be adjusted accordingly using serial dilutions. Viability should then be estimated by adding Trypan blue (0.4%) to zoospores (1:2 dilution) and used to differentiate live and dead cells (Stockwell et al. 2010). Flow cytometry is another technique that can be used to count zoospores and estimate viability, although a protocol for *Bsal* zoospores has not yet been developed.

## 5. Isolation of *Bsal* from infected tissue

*Bsal* can be isolated from infected host tissue using methods described by Martel et al. (2013) for *Bsal* as well as by methods developed for *Bd* (Longcore et al. 1999, Waddle et al. 2018, Fisher et al. 2018, Cook et al. 2018). Tissues used for culturing should be collected from areas with prevalent or suspected *Bsal* colonization such as ulcers or highly keratinized toe tips. Ideally tissues should be collected from an infected animal directly after euthanasia before tissues are overrun with bacterial growth. If possible, collect several tissue samples from each animal to maximize the likelihood of successful isolation. Tissues should be dragged through TGhL agar plates containing 200 mg/L penicillin and 200 mg/L streptomycin antibiotics to remove any bacteria or other fungal species which are present. The clean tissue can now be placed on a new TGhL agar plate and incubated at 15 C. Observe plates daily and discard contaminated plates. Once motile zoospores are observed (3 to 6 days), the tissue can be carefully removed from the TGhL plate. The *Bsal* zoospores and zoosporangia can then be removed using a sterile 25 cm cell scraper and transferred to a flask of liquid media. Each culture should be monitored closely thereafter for signs of contamination.

## 6. Cryopreservation

### 6.1 Freezing

For long term cryopreservation of *Bsal*, we have adopted methods developed for *Bd* (Boyle et al. 2003, Gleason et al. 2007). Because cell health at the time of freezing is critical, start with a liquid culture that is actively growing and has plenty of motile zoospores. While we only provide recommendations for cryopreserving mixed cultures, synchronized zoospores have also been successfully revived using this approach. Cryopreservation in −80 C freezers can last for months, and we have recovered *Bd* and *Bsal* cultures after storage in liquid nitrogen for years.

First, remove zoospores and sporangia from the culture flask. Because sporangia tightly adhere to the walls of TC treated flasks, use a sterile cell scraper to dislodge sporangia from the flask walls. Transfer the culture to a sterile tube and centrifuge at 2500 rcf for 5 minutes. To prevent the pelleted cells from swimming back into solution, pour off and discard the supernatant immediately after the centrifuge stops. Resuspend the cell pellet in full strength TGhL culture media supplemented with 10% sterile dimethylsulfoxide (DMSO) pre-chilled to 4 C.

Aliquot 1 mL of the cell solution into pre-labeled cryovials equipped with gaskets and transfer to an isopropanol container (e.g. “Mr. Frosty” Sigma cat.# C1562 or equivalent, note: this is essential) that has been pre-chilled to −80 C. Place the isopropanol container in a −80 C freezer for 24 to 48 hours. Move the frozen cryovials into liquid nitrogen for long-term storage.

### 6.2 Thawing

To thaw, remove one cryovial from liquid nitrogen and swirl in a beaker with lukewarm water (approximately 25 to 35 C), until the contents are just thawed, being careful to keep the lip of the sealed tube above the waterline. Add the thawed sample to 10 mL of culture media pre-chilled to 15 C. Using additional media, rinse any remaining cells from the cryovial and combine. To remove the DMSO, centrifuge the media-cell mixture at 2500 rcf for 5 minutes, immediately discard the supernatant, and resuspend the cell pellet in 10 mL of culture media pre-chilled to 15 C. Transfer the cells to a 25 mm^2^ TC treated flask and examine using a inverted microscope. If the cells are confluent after settling to the bottom of the flask, we recommend diluting an aliquot 1:10 into an additional flask and monitoring both to make sure at least one is not overcrowded. Incubate the culture(s) at 15 C and observe daily. When swimming zoospores are visible, the culture is ready to begin liquid subculturing (see ‘Subculturing in liquid media’).

## 7. Conclusion

The amphibian chytrid pathogen *Bsal* poses a significant threat to global biodiversity. The ability to develop the mitigation and conservation strategies required to effectively manage a *Bsal* outbreak is limited by large gaps in our knowledge about the basic biology of this deadly pathogen. We provide recommendations for growth and handling to help close these gaps by lowering the barriers for new researchers working with *Bsal*, and by facilitating the comparison of results from different laboratories. We therefore recommend that researchers follow the procedures outlined here and describe any necessary deviations from these procedures in as much detail as is practical in the materials and methods section of the relevant manuscript.

## 8. Recipes

### 8.1 Liquid media

After adding each ingredient listed below, bring the volume up to 1 liter (L) with deionized water (diH_2_O). Autoclave all mixtures before use and/or storage. Liquid media can be stored at 4 C, 15 C, or room temperature but should be brought to 15 C prior to being used for culturing.

#### Full-strength TGhL (1L)

16 g tryptone

4 g gelatin hydrolysate

2 g lactose

#### Half-strength TGhL (1L)

8 g tryptone

2 g gelatin hydrolysate

1 g lactose

#### Full-strength 1% Tryptone (1L)

10 g tryptone

#### Half-strength 1% Tryptone (1L)

5 g tryptone

#### mPmTG (1L)

0.5 g peptonized milk

0.5 g tryptone

2.5 g glucose

#### Provosoli at pH 7.0 (1L)

Add 2 mL of each salt:

NaNO_3_ (6.25 g in 250 mL diH_2_O)

MgSO_4_ · 7H_2_O (5.0 g in 250 mL diH_2_O)

CaCl_2_ · 2H_2_O (3.3 g in 250 mL diH_2_O)

K_2_HPO_4_ (0.75g in 250 mL diH_2_O)

KCl (6.25g in 250 mL diH_2_O)

KH_2_PO_4_ (0.75 g in 250 mL diH_2_O)

#### Bonner’s Salts (1L)

0.6 g NaCl

0.75 g KCl

0.3 g CaCl_2_

### 8.2 Solid media

Prepare TGhL (full or half-strength) following the recipes for liquid media. Add 10 g of biological grade agar per 1L of liquid media. Autoclave. In general, 250 mL of the agar-TGhL media generates approximately 20 plates (60 mm). Plates can be stored at 4 C for several months but should be brought to 15 C prior to being used for culturing.

If using antibiotics, it is important that autoclaved agar-TGhL media is first cooled to 60 C. Even cooling can be achieved within a water-bath set to 45 C to 60 C.

### 8.3 Antibiotics

#### Stock antibiotics for liquid media

Individually, combine each antibiotic with 5 mL nanopure water:

0.5 g Ampicillin sodium salt (371.39 g/mol)

0.625 g Streptomycin sulfate (1,457.39 g/mol)

Because antibiotics are destroyed by autoclaving, use a sterile 0.2 µm cellulose acetate syringe to filter each antibiotic into a sterile tube to sterilize. Aliquot sterilized stock mixtures and store at −20 C. Prior to using, thaw aliquots at room temperature and use a sterile pipet to add 10 µL of each stock mixture per 10 mL liquid media.

#### Antibiotics for solid media

Dissolve 0.1 g of Ampicillin sodium salt (371.39 g/mol) and 0.1 g of Streptomycin sulfate (1,457.39 g/mol) in 2 mL nanopure water. Sterilize by filtering through a 0.2 µm cellulose acetate syringe filter. Add 1 mL prepared antibiotics per 500 mL autoclaved solid media that has been cooled to 60 C. Gently swirl to mix and pour plates as normal.

## 9. Acknowledgements

We would like to thank Amanda Tokash-Peters for data collection on prelimary growth assays and Dr. Louise Rollins-Smith for helpful discussions. This work was supported by the National Science Foundation (IOS-1827257 to L.F.-L. and 1814520 to M.J.G.).

## References

Absher M (1973) Hemocytometer. Tissue Culture:395–397

Berger L, Speare R, Daszak P, Green DE, Cunningham AA, Goggin CL, Slocombe R, Ragan MA, Hyatt AD, McDonald KR, Hines HB, Lips KR, Marantelli G, Parkes H (1998) Chytridiomycosis causes amphibian mortality associated with population declines in the rain forests of Australia and Central America. Proc Natl Acad Sci USA 95:9031–9036

Boyle DG, Hyatt AD, Daszak P, Berger L, Longcore JE, Porter D, Hengstberger SG, Olsen V (2003) Cryo-archiving of Batrachochytrium dendrobatidis and other chytridiomycetes. Dis Aquat Organ 56:59–64

Burnett LC, Lunn G, Coico R (2009) Biosafety: guidelines for working with pathogenic and infectious microorganisms. Curr Protoc Microbiol Chapter 1:Unit 1A.1–1A.1.14

Chatfield MWH, Moler P, Richards-Zawacki CL (2012) The amphibian chytrid fungus, Batrachochytrium dendrobatidis, in fully aquatic salamanders from Southeastern North America. (J Sturtevant, Ed.). PLoS ONE 7:e44821

Cook KJ, Voyles J, Kenny HV, Pope KL, Piovia-Scott J (2018) Non-lethal isolation of the fungal pathogen Batrachochytrium dendrobatidis (Bd) from amphibians. Dis Aquat Organ 129:159–164

Coté RJ (2001) Aseptic technique for cell culture. (JS Bonifacino, M Dasso, JB Harford, J Lippincott-Schwartz, and KM Yamada, Eds.). Curr Protoc Cell Biol Chapter 1:Unit 1.3–1.3.10

Fisher MC, Ghosh P, Shelton JMG, Bates K, Brookes L, Wierzbicki C, Rosa GM, Farrer RA, Aanensen DM, Alvarado-Rybak M, Bataille A, Berger L, Böll S, Bosch J, Clare FC, A Courtois E, Crottini A, Cunningham AA, Doherty-Bone TM, Gebresenbet F, Gower DJ, Höglund J, James TY, Jenkinson TS, Kosch TA, Lambertini C, Laurila A, Lin C-F, Loyau A, Martel A, Meurling S, Miaud C, Minting P, Ndriantsoa S, O’Hanlon SJ, Pasmans F, Rakotonanahary T, Rabemananjara FCE, Ribeiro LP, Schmeller DS, Schmidt BR, Skerratt L, Smith F, Soto-Azat C, Tessa G, Toledo LF, Valenzuela-Sánchez A, Verster R, Vörös J, Waldman B, Webb RJ, Weldon C, Wombwell E, Zamudio KR, Longcore JE, Garner TWJ (2018) Development and worldwide use of non-lethal, and minimal population-level impact, protocols for the isolation of amphibian chytrid fungi. Sci Rep 8:7772–8

Fritz-Laylin LK, Lord SJ, Mullins RD (2017) WASP and SCAR are evolutionarily conserved in actin-filled pseudopod-based motility. The Journal of Cell Biology 216:1673–1688

Garner TWJ, Schmidt BR, Martel A, Pasmans F, Muths E, Cunningham AA, Weldon C, Fisher MC, Bosch J (2016) Mitigating amphibian chytridiomycoses in nature. Philos Trans R Soc Lond, B, Biol Sci 371:20160207

Gleason FH, Mozley-Standridge SE, Porter D, Boyle DG, Hyatt AD (2007) Preservation of Chytridiomycota in culture collections. Mycological Research 111:129–136

Gower DJ, Doherty-Bone T, Loader SP, Wilkinson M, Kouete MT, Tapley B, Orton F, Daniel OZ, Wynne F, Flach E, Müller H, Menegon M, Stephen I, Browne RK, Fisher MC, Cunningham AA, Garner TWJ (2013) Batrachochytrium dendrobatidis infection and lethal chytridiomycosis in caecilian amphibians (Gymnophiona). Ecohealth 10:173–183

Kilpatrick AM, Briggs CJ, Daszak P (2010) The ecology and impact of chytridiomycosis: an emerging disease of amphibians. Trends Ecol Evol (Amst) 25:109–118

Laking AE, Ngo HN, Pasmans F, Martel A, Nguyen TT (2017) Batrachochytrium salamandrivorans is the predominant chytrid fungus in Vietnamese salamanders. Sci Rep 7:44443

Langhammer PF, Lips KR, Burrowes PA, Tunstall T, Palmer CM, Collins JP (2013) A fungal pathogen of amphibians, Batrachochytrium dendrobatidis, attenuates in pathogenicity with in vitro passages. (MMC Fisher, Ed.). PLoS ONE 8:e77630

Lips KR (2016) Overview of chytrid emergence and impacts on amphibians. Philos Trans R Soc Lond, B, Biol Sci 371:20150465

Longcore JE, Pessier AP, Nichols DK (1999) Batrachochytrium Dendrobatidis gen. et sp. nov., a Chytrid Pathogenic to Amphibians. Mycologia 91:219–227

Martel A, Spitzen-van der Sluijs A, Blooi M, Bert W, Ducatelle R, Fisher MC, Woeltjes A, Bosman W, Chiers K, Bossuyt F, Pasmans F (2013) Batrachochytrium salamandrivorans sp. nov. causes lethal chytridiomycosis in amphibians. Proc Natl Acad Sci USA 110:15325–15329

Morgan JAT, Vredenburg VT, Rachowicz LJ, Knapp RA, Stice MJ, Tunstall T, Bingham RE, Parker JM, Longcore JE, Moritz C, Briggs CJ, Taylor JW (2007) Population genetics of the frog-killing fungus Batrachochytrium dendrobatidis. Proc Natl Acad Sci USA 104:13845–13850

Refsnider JM, Poorten TJ, Langhammer PF, Burrowes PA, Rosenblum EB (2015) Genomic Correlates of Virulence Attenuation in the Deadly Amphibian Chytrid Fungus, Batrachochytrium dendrobatidis. G3 (Bethesda) 5:2291–2298

Sabino-Pinto J, Veith M, Vences M, Steinfartz S (2018) Asymptomatic infection of the fungal pathogen Batrachochytrium salamandrivorans in captivity. Sci Rep 8:11767–8

Scheele BC, Pasmans F, Skerratt LF, Berger L, Martel A, Beukema W, Acevedo AA, Burrowes PA, Carvalho T, Catenazzi A, la Riva De I, Fisher MC, Flechas SV, Foster CN, Frías-Álvarez P, Garner TWJ, Gratwicke B, Guayasamin JM, Hirschfeld M, Kolby JE, Kosch TA, La Marca E, Lindenmayer DB, Lips KR, Longo AV, Maneyro R, McDonald CA, Mendelson J, Palacios-Rodriguez P, Parra Olea G, Richards-Zawacki CL, Rödel M-O, Rovito SM, Soto-Azat C, Toledo LF, Voyles J, Weldon C, Whitfield SM, Wilkinson M, Zamudio KR, Canessa S (2019) Amphibian fungal panzootic causes catastrophic and ongoing loss of biodiversity. Science 363:1459–1463

Schloegel LM, Toledo LF, Longcore JE, Greenspan SE, Vieira CA, Lee M, Zhao S, Wangen C, Ferreira CM, Hipolito M, Davies AJ, Cuomo CA, Daszak P, James TY (2012) Novel, panzootic and hybrid genotypes of amphibian chytridiomycosis associated with the bullfrog trade. Mol Ecol 21:5162–5177

Stegen G, Pasmans F, Schmidt BR, Rouffaer LO, Van Praet S, Schaub M, Canessa S, Laudelout A, Kinet T, Adriaensen C, Haesebrouck F, Bert W, Bossuyt F, Martel A (2017) Drivers of salamander extirpation mediated by Batrachochytrium salamandrivorans. Nature 544:353–356

Stockwell MP, Clulow J, Mahony MJ (2010) Efficacy of SYBR 14/propidium iodide viability stain for the amphibian chytrid fungus Batrachochytrium dendrobatidis. Dis Aquat Organ 88:177–181

Van Rooij P, Martel A, Haesebrouck F, Pasmans F (2015) Amphibian chytridiomycosis: a review with focus on fungus-host interactions. Vet Res 46:137–22

Van Rooij P, Pasmans F, Coen Y, Martel A (2017) Efficacy of chemical disinfectants for the containment of the salamander chytrid fungus Batrachochytrium salamandrivorans. (J Kerby, Ed.). PLoS ONE 12:e0186269

Waddle AW, Sai M, Levy JE, Rezaei G, van Breukelen F, Jaeger JR (2018) Systematic approach to isolating Batrachochytrium dendrobatidis. Dis Aquat Organ 127:243–247

Watts A, Olson D, Harris R, Mandica M (2019) The deadly amphibian bsal disease: How science-management partnerships are forestalling amphibian biodiversity losses. Science Findings 214

Woodhams DC, Alford RA (2005) Ecology of chytridiomycosis in rainforest stream frog assemblages of tropical Queensland. Conservation Biology 19:1449–1459

Yuan Z, Martel A, Wu J, Van Praet S, Canessa S, Pasmans F (2018) Widespread occurrence of an emerging fungal pathogen in heavily traded Chinese urodelan species. Conservation Letters 11

Batrachochytrium: Biology and Management of Amphibian Chytridiomycosis Batrachochytrium: Biology and Management of Amphibian Chytridiomycosis.: 1–18

Science Findings: Pacific Northwest Research Station Science Findings: Pacific Northwest Research Station.

